# MAJIQ V3 offers improvements in accuracy, performance, and usability for splicing analysis from RNA sequencing

**DOI:** 10.1101/2024.07.02.601792

**Authors:** Joseph K Aicher, Barry Slaff, San Jewell, Yoseph Barash

## Abstract

**Summary:** RNA splicing plays important roles in cell-type-specific expression of gene isoforms and is causally linked to numerous diseases. Previously, our group developed MAJIQ V1 and V2 for RNA splicing detection, quantification, and visualization from RNA sequencing. Here we report an update, MAJIQ V3, that offers several improvements in time, memory, accuracy, and usability. We demonstrate the benefits of MAJIQ V3 using GTEx data for different analysis scenarios involving different dataset sizes.

## Introduction

RNA alternative splicing (AS) is a process by which different subsets of pre-mRNA segments are jointed, or spliced, together. AS plays important roles in cell-type-specific expression of gene isoforms. Up to 95% of human multi-exon genes are alternatively spliced (AS) [1, 2], and changes to splicing are causally linked to numerous diseases[3, 4]. The importance of RNA splicing has led to the development of computational tools which discover, quantify, and compare RNA splicing variations between groups of RNA sequencing experiments[5]. Previously, our group developed Modeling Alternative Junction Inclusion Quantification (MAJIQ) V1 and V2 for RNA splicing detection, quantification, and visualization from RNA sequencing [6, 7]. Most recently, MAJIQ was extended to integrate long reads and full isoforms analysis [8].

Here we report MAJIQ V3, an update which improves upon the performance and features offered in previous versions. The MAJIQ V3 components are the Builder, MOCCASIN, Quantifier, and VOILA (Fig 1a). The MAJIQ Builder detects AS events from RNA sequencing experiments and quantifies read coverage for the detected events. These AS events are defined over each gene’s splice graph as local splicing variations (LSV), in which multiple splice junctions or intron retention (IR) are coming out or into a given exon (see illustration in Fig 1b). MOCCASIN[9] is a method to adjust read coverage for known and unknown confounding factors in the detected LSVs quantifications which was previously provided as a separate program. The MAJIQ Quantifier uses the (optionally MOCCASIN-adjusted) LSV read coverage to estimate a beta-distributed posterior for inclusion (PSI) for each splice junction or intron within each LSV. As in previous versions, the beta posterior considers the total evidence (coverage) for PSI as well as the distribution of reads across genomic positions, with low-coverage LSVs having higher variance. The Quantifier also includes algorithms to calculate differential inclusion between two groups of replicate experiments (MAJIQ DeltaPSI) and non-replicate (heterogeneous) experiments (MAJIQ-HET). VOILA provides graphical visualizations of quantified splicing events and includes a Modulizer for identifying different patterns of high-coverage splicing variations.

Compared to MAJIQ V2, V3 includes performance enhancements driven by a more efficient reimplementation with contiguous data structures; a finer-grained command-line interface (CLI) allowing for more user control, incremental addition of new experiments to previous analyses, and parallelized coverage calculations; and improved quantification algorithms. In the following sections, we summarize the key improvements and demonstrate the benefits using GTEx data[10] for different analysis scenarios involving small to large datasets.

### Decoupled build steps and new splicegraph operations

MAJIQ V2 and V3 parse aligned reads from BAM files into “SJ” files with coverage over junctions and retained introns. These SJ files are used as input rather than BAM files to the MAJIQ build step. In the build step, SJ files are organized into independent build groups which, along with GFF3 transcriptome annotations, are processed all at once in the MAJIQ V2 builder to simultaneously produce a splice graph (collection of LSVs) per gene, as well as coverage over these LSVs. In contrast, MAJIQ V3 decouples these two elements into separate commands. As a result, coverage over LSVs can be produced directly from an input splicegraph and SJ coverage (Fig 1c). This improves parallelization of splicing analysis and allows incremental quantification of new experiments for direct comparison with previous analyses. MAJIQ V3 additionally introduces the following options:

1. *Combine splicegraphs*. New splicegraphs can be generated incrementally by combining two or more input splicegraphs (Fig 1d). The combined splicegraph is structurally equivalent to a splicegraph built with the combined experiments used to create the input splicegraphs. Please see Supplemental Methods for details of the logic to match junctions, introns, and LSVs between splicegraphs.
2. *Generate PSI coverage for one splicegraph, excluding all LSVs in another splicegraph*. When incrementally adding new samples to a previous analysis, this can be used to quantify psi coverage for only the new LSVs rather than re-quantifying all LSVs for all samples, the approach needed in previous MAJIQ versions.
3. *Identify novel junctions, retained introns, and LSVs relative to another splicegraph*. This ability can be used for the clinical setting to identify transcriptomic features which are unique to a patient relative to one or more controls. This differs from MA-JIQ V2, which always defines “de novo” introns and junctions relative to annotated transcripts, not other experiments. Please see Supplemental Methods for the details of identifying matched junctions, introns, and LSVs.

### Improvements to Retained Intron quantifications

MAJIQ V2 calculates junction and intron coverage for each gene independently using reads that overlap the relevant region. However, since genes can overlap in genomic coordinates, this approach suffers from two issues. First, there is redundancy in coverage quantification for overlapping regions. Second, counts for an exon of one gene may be counted towards introns of another gene. In contrast, MAJIQ V3 measures coverage over non-overlapping regions (broken by strand if the RNA-seq experiment is stranded). The intronic regions from each gene are combined, and the exonic regions for each region are subtracted, as illustrated in Figure 1e. This excludes any exonic region from contributing to intronic coverage in a different gene. Additional care is taken to identify intronic regions that belong to “annotated introns”. Annotated introns are part of some annotated transcript’s exon that was split by junctions from other transcripts. We identify regions that correspond to annotated introns so that they are not counted to novel introns. MAJIQ V3 measures coverage over these regions using the same procedure that MAJIQ V2 uses for gene introns[7].

**Figure 1.**
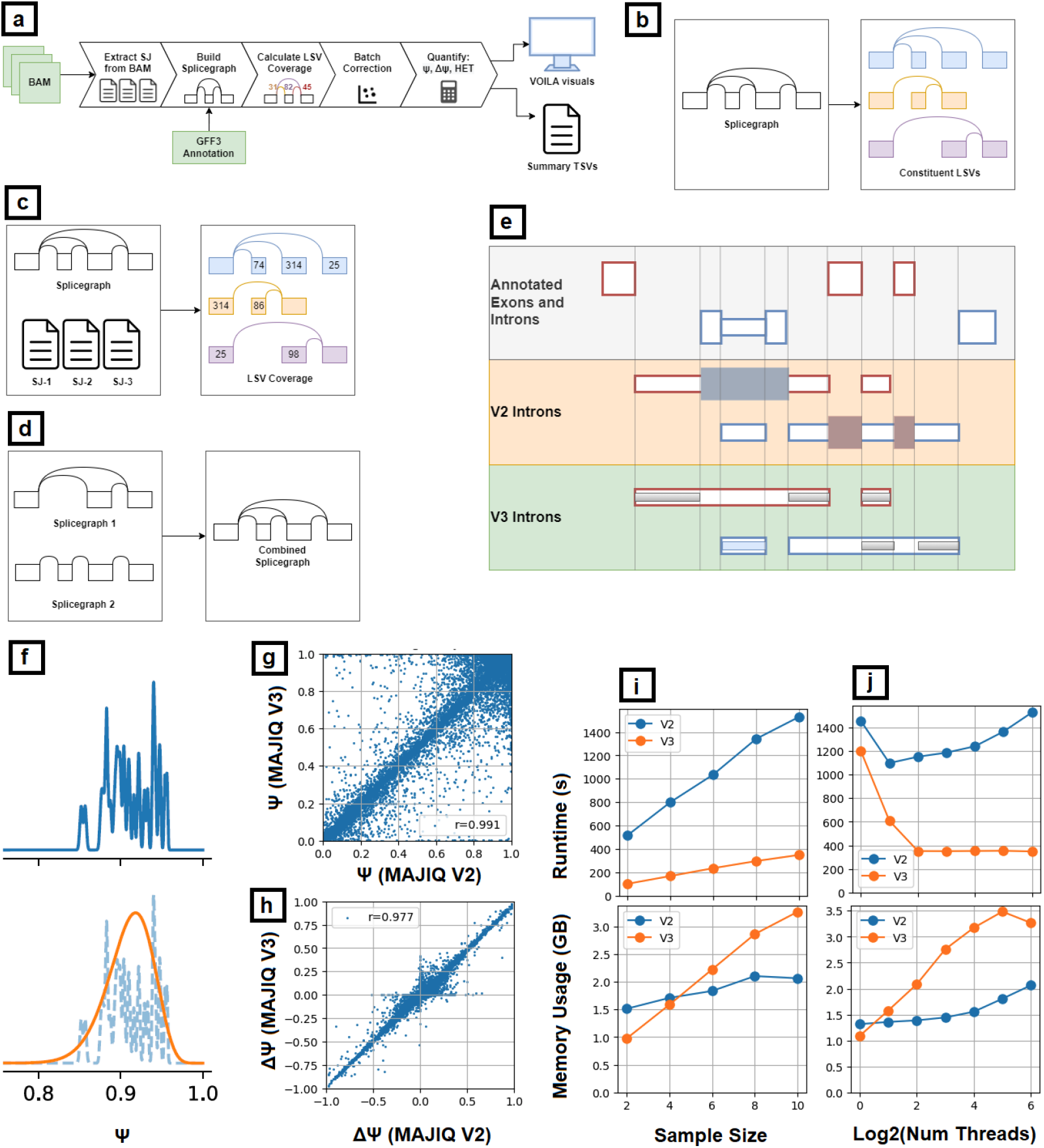
MAJIQ V3 analysis pipeline and improvements over MAJIQ V2. (a) The MAJIQ V3 splicing analysis pipeline. RNA-seq BAM files are extracted to “SJ” files with read counts for splice junctions and introns; SJ files are combined with a GFF3 genome annotation to build a splicegraph (SG) with all the local splicing variations (LSVs). Next, read coverage over LSVs is quantified and MOCCASIN can be optionally applied for batch correction. Finally, the MAJIQ Quantifier estimates inclusion (PSI) and/or differential inclusion (DeltaPSI, HET) for LSVs splice junctions and introns. (b) A splicegraph (left) with exons (rectangles) and junctions (curved lines) broken into its three constituent LSVs (right). LSVs are characterized by a source or target exon; here, the top LSV (blue) is a source-exon LSV and the bottom two (orange, purple) are target-exon LSVs. (c) In MAJIQ V3, coverage over LSVs can be computed directly from an input splicegraph and SJ coverage files. This improves parallelization of splicing analysis and allows incremental quantification of new experiments for direct comparison with previous analyses. (d) In MAJIQ V3, new splicegraphs can be generated incrementally by combining 2 or more input splicegraphs. The combined splicegraph is structurally equivalent to a splicegraph built with the combined experiments used to create the input splicegraphs. (e) Diagram illustrating how MAJIQ V2 and V3 measure coverage in overlapping regions which can overlap with each others’ exons. For V2, oversized dark rectangles show where exonic coverage from other genes might lead to artifactual coverage. MAJIQ V3 instead measures coverage over non-overlapping regions which exclude all exons. Coverage regions (thin interior rectangles) are defined by excluding all exonic regions and noting which regions originated from annotated introns. Coverage over introns is determined by averaging overlapping regions. Coverage over the interval of an annotated retained intron (blue coverage region) counts only towards that intron and not possible novel introns. (f) MAJIQ V3 uses a smooth approximation to the uniform mixture of bootstrapped posteriors. Shown is a smooth approximation (orange) of a mixture (blue) of 30 beta distributions. This improves the computational speed and stability of sampling from posterior PSI and calculating quantiles. (g) MAJIQ V3 vs MAJIQ V2 PSI comparison using 10 cerebellum RNA-seq experiments from GTEx. (h) MAJIQ V3 vs MAJIQ V2 delta-PSI comparison for 10 cerebellum vs 10 muscle-skeletal GTEX samples. (i) Runtime and peak memory vs group size for the build and LSV quantification steps for the small evaluation set (10 vs 10 samples). (j) Runtime and peak memory vs thread count for the build and LSV quantification steps for the small evaluation set (10 vs 10 samples).

### Improved PSI posterior calculations

MAJIQ V3 improves the estimation of variance in PSI posterior distribution. For every junction in every LSV, MAJIQ repeats M times bootstrap sampling of reads to create M beta posterior distributions for Ψ. MAJIQ V2 calculates variance by discretizing Ψ into 40 equally-sized bins over [0, 1], and setting the probability of each bin to the average probability of the bin over the M beta distributions. Then, MAJIQ V2 calculates variance with respect to the midpoints of the bins, weighted by these probabilities. This procedure can be thought of as an approximation of a uniform mixture of beta distributions. Thus, the variance of Ψ is now calculated instead in V3 using the law of total variance applied to mixture of Ψ distributions, making the calculation both faster and more accurate (for details, please see Supplemental Methods).

Another change introduced in V3 is the usage of a smooth approximation to the mixture of bootstrapped PSI posteriors. This change addresses challenges with modeling PSI directly as a finite mixture distribution as total coverage at an LSV increases. As total coverage increases, the uncertainty of each mixture component becomes vanishingly small. Since we only sample a finite number of mixture components, the support of the actual mixture distribution becomes finite. This leads to negative consequences when sampling or taking quantiles from the distribution. To resolve such potential issues, MAJIQ approximates the mixture distribution with a single beta distribution, with parameters set to match the mixture distribution’s mean and variance (Fig 1f), with variance calculated as described above.

### MOCCASIN batch correction improvements

MOCCASIN is a method to adjust junction and intron coverage for known and unknown confounding factors. The user can optionally input known confounders and/or ask MOCCASIN to discover unknown confounders. Then, MOCCASIN calculates and outputs adjusted coverage[9]. Instead of a separate package, MAJIQ V3 now includes a rewritten implementation of MOCCASIN. The new implementation is vectorized over junctions and bootstrap replicates, resulting in significant performance improvements. It also takes advantage of changes to underlying data structures to remove no longer necessary steps (e.g., matching indexes for events, which are now consistently ordered) and to enable multithreaded and/or distributed parallelism using Dask.

As part of MOCCASIN’s new implementation, MAJIQ V3 decouples the MOCCASIN steps to support incrementally applying the algorithm to new samples and/or LSVs. The newly decoupled steps include:

1. *factors-model*: Build a model of unknown confounders.
2. *factors-infer* : Augment the input factors with inferred unknown confounders.
3. *coverage-model*: Build a model of coverage vs. confounding factors.
4. *coverage-infer* : Calculate and output adjusted coverage over LSVs.

### Updated data structures

MAJIQ V3 introduces several changes to the underlying data structures to improve time and memory efficiency. Specifically, splicegraph, coverage and VOILA files were rewritten to improve performance, decrease file sizes, and enable parallelization with Dask. Redundant information (i.e. gene names, contig names) is stored either by reference or computed as needed to decrease storage and memory usage. For SJ coverage, per-bin read counts are now kept in storage as a sparse matrix (most junctions only have a few nonzero bins/positions). Bootstrapping is no longer stored and instead computed on the fly as needed. This means that SJ coverage is now deterministic/reproducible for the same input BAM file on different runs of MAJIQ. Splicegraphs exclude read counts, which are split into separate splicegraph coverage files (produced by the sg-coverage command). This keeps splicegraphs relatively small and saves time by assessing coverage when requested rather than automatically for all input samples. Coverage over LSVs is stored redundantly, storing the percentage of reads in the LSV to which a junction or intron was assigned in one array and the total number of reads in the LSV repeated for each junction and intron. This is to enable chunking the data over junctions/retained introns when the number of samples is large such that it can no longer be loaded all at once in memory. LSVs are always stored in the same order to maintain efficiency.

### VOILA data storage and retrieval optimizations

VOILA is the visualization package for MAJIQ. For MAJIQ V3 compatibility and improved performance the VOILA underlying system for data storage and retrieval has been overhauled. In the V2 version of VOILA, a sqlite database, HDF5 matrix files, and python pickle format files were read in tandem to process and display data. The usage of HDF5 matrix files in particular were a bottleneck in terms of read/write performance, as the paradigm involved accessing many small sub-matrices for each LSV, which caused a bottleneck, especially when accessed in parallel and over a network file system, as they did not behave well with usual Linux NFS cache mechanisms. Both of these use-cases are common, so their replacement in V3 was a necessary performance improvement.

For the V3 version of VOILA, the sqlite and HDF5 files were replaced with Zarr type datasets, which are more performant when reading and writing in parallel and over network file systems. This was achieved not only by taking advantage of built-in parallelism of the zarr format, but also re-structuring the codebase to use fewer, large matrices instead of a multitude of smaller ones. The Python pickle files are still retained for some auxiliary analysis types, e.g. small dataset sizes which fit entirely in memory.

### Performance Evaluations

We demonstrate the benefits of MAJIQ V3 using GTEx[10] data for different analysis scenarios involving small to large datasets. Evaluations were performed on a TS1-160859871 server with 64 core AMD EPYC Milan 7713 CPUs, 2TB of RAM, and 100GB network attached SSD storage. We use two subsets of GTEx data: a “small” analysis consisting of up to 10 samples from each of two tissue types (20 samples total), and a “large” analysis consisting of 10 samples from each of 20 tissue types (200 samples total). First, Fig 1g-h demonstrates that MAJIQ V2 and V3 quantifications are largely the same, with LSV deviating more than 10% in only 3.6% of PSI and 1.6% of dPSI cases. This small set of LSV with larger deviations are due to V3 improvements such as better retained-intron quantifications described above.

Next, we illustrate the performance benefits of MAJIQ V3. When using the smaller dataset of up to 20 samples V3 runs significantly faster than V2. These observations hold while increasing the number of samples per group from 2 to 10 (Fig 1i). The peak memory footprint of MAJIQ V3 in these runs is 3.5 GB, well within the capabilities of modern laptop and desktop computers. For the larger data involving 200 samples, MAJIQ V3 demonstrates a significant speedup when multiple threads are used. For example, V3 completes the build and all 190 pairwise comparative quantifications across the 20 tissue groups (MAJIQ DeltaPSI and MAJIQ-HET) in only 126 minutes using 64 threads, which is 3.2x faster than to MAJIQ V2. In the same runs, the memory peak of MAJIQ V3 was 11.8 GB, compared to the 4.2 GB peak of MAJIQ V2. Notably, in both V2 and V3 versions, the build step has both the highest memory peak and longest runtime. While V3 consumes more memory in this multi-threaded execution, we note that 11.8 GB is well within the capabilities of modern laptop and desktop computers intended to do data analytics of large data with hundreds of samples. Overall, these results illustrate the capability of V3 to quickly perform analyses with large and heterogeneous data while leaving a modest memory footprint.

### Installation and Usage

MAJIQ V3 is implemented in Python and C++. We recommend installing MAJIQ V3 using pip or Conda and invoking MAJIQ commands from the terminal. We recommend Python version 3.10 and Ubuntu 22.04 or above for the smoothest experience, however, it is possible to install and use MAJIQ on other Linux, macOS, and even Windows if the correct system libraries are available.

In order to install MAJIQ, one should clone and install the MOCCASIN repository, followed by the MAJIQ repository after agreeing to the BioCiphers license agreement on the Biociphers website. Following this, one may use python venv or Conda vent to make a new python3.10 virtual environment, activate it, and install MOCCASIN with pip, followed by MAJIQ.

There may be some system libraries required, such as HTSlib, zlib, etc. Pip/Conda will show a message during the installation if any common libraries are missing prior to installation.

MAJIQ requires RNA sequencing experiments input as samtools-indexed BAM files and an genome annotation input as a GFF3 file. The MAJIQ builder extracts each BAM file into a new file including reads over splice junctions and introns (MAJIQ SJ format). Since the downstream builder steps depend only on the SJ file for each experiment, the BAM extraction step need only take place once per experiment.

## Availability and Implementation

MAJIQ V3 is implemented in Python and C++. Please check majiq.biociphers.org for more details, documentation, and exact release date.

## Supplemental MAJIQ V3 Methods

### Detecting Strandedness

MAJIQ V3 automatically detects the strandedness of the input RNA-seq experiments if not provided by the user. At a high level, if all the reads mapping to a junction are in one direction, either the experiment is stranded or we are seeing a rare event (depending on the total number of reads) from an unstranded case. If an experiment is stranded, we expect to see a lot of reads matching our annotated junctions in the directions associated with the strandedness of the experiment. We look at all junctions to see if we see the same kind of bias across junctions. The algorithm is:

1. MAJIQ calculates the number of aligned reads per junction *j* with original *o* and reverse *r* strandedness assumed: *n*_*j,o*_ and *n*_*j,r*_.
2. MAJIQ restricts the analysis to junctions having a min-threshold number *T* of total aligned reads: *n*_*j,o*_ + *n*_*j,r*_ > *T*.
3. MAJIQ calculates the ratio per junction *r*_*j*_ of aligned reads counts for original strandedness and both: *r*_*j*_ = *n*_*j,o*_/(*n*_*j,o*_ + *n*_*j,r*_).
4. MAJIQ calculates junctions: *r*_med_ = median(*r*_*j*_)_*j*∈*J*_
5. If the median ratio deviates from 0.5 by more than min-deviation (0.2), strandedness is called; otherwise the experiments are assumed unstranded.

### Retained-Intron Event Acceptance Model

At a high level, we expect the readrate to be near-constant across all nucleotide positions of a retained intron; large deviations would indicate that the candidate intron is not really one retained intron. MAJIQ V3 determines thresholds for acceptance of MAJIQ introns of various lengths by matching junction acceptance threshold probability with identical per-position readrate.

The model is that each position on a transcript feature (junction, intron) has the same underlying readrate. We observe reads for each position as a Poisson random variable with this underlying readrate.

We accept junctions on the condition that they have at least minreads reads observed across all bins and minpos bins with at least 1 read. For some acceptance probability q, we use Newton-Raphson iteration to backsolve for minimum readrate required to pass thresholds with that probability.

Then, we can backsolve the intron thresholds for each intron length L that pass with just under the same probability q when given equivalent per-position readrate.

### Identifying matched junctions, introns, and LSVs

These operations were motivated by two goals we had for the clinical pipeline. First, we wanted MAJIQ to identify novel junctions and retained introns relative to control RNA-seq experiments (as well as annotated transcripts). In contrast, MAJIQ V2 always defines “de novo” status of introns and junctions relative to annotated transcripts. Second, when new samples (or cases) are added to an analysis (of controls), the existing samples have already been quantified with respect to the old set of LSVs. We also expect that most of these LSVs are unchanged in the new splicegraph. Rather than redundantly storing and requantifying these same LSVs, identifying matched LSVs between the new and old splicegraphs would permit producing coverage for (and subsequently quantifying) only the LSVs unique to the new splicegraph.

Identifying matched junctions is straightforward: does the other splicegraph have a junction for the same gene and coordinates. Identifying matched retained introns requires more care. Exonic, and thus intronic coordinates, are updated between splicegraphs to match novel junction splice sites. So, we cannot simply match intron coordinates. It is not enough to look for overlapping intron coordinates, either. Extended exon boundaries and novel exons can mask or split introns. So, we match introns from both splicegraphs to regions corresponding to boundaries of annotated exons (as done for combining splicegraphs). Two introns are matched if they share the same annotated intronic region. Finally, MAJIQ identifies matched events by searching for events belonging to the same gene which have junctions and introns with the same coordinates. In this case, we require intron coordinates to be identical because matched events are most useful for thinking about quantifications. Different intron coordinates could lead to different intron coverage and thus quantification.

Using these operations, MAJIQ v3 introduces the optional flag –annotated to its quantification commands to identify novel transcriptomic features relative to another splicegraph. When used, MAJIQ accepts an additional splicegraph as input and uses the above operations to identify which junctions, retained introns, and events are novel relative to this splicegraph. Otherwise, MAJIQ V3 has the same behavior as MAJIQ V2 and marks junctions and retained introns with de novo status relative to the annotated transcripts.

### Corrected posterior standard deviation / variance

We correct the calculation of PSI posterior variance used in previous MAJIQ versions. MAJIQ estimates *M* bootstrap replicates of beta posterior distributions for Ψ. MAJIQ V2 calculates variance by discretizing Ψ into 40 equally-sized bins over [0, 1], and setting the probability of each bin to the average probability of the bin over bootstrap replicates. If we imagine increasing the number of equally-sized bins, we see that this is an approximation of a uniform mixture of beta distributions. The variance of this distribution can be directly and exactly calculated much more efficiently.

This mixture distribution can be factored into its mixture components using the hidden random variable *Z*∼ Uniform(1, …, *M*), indicating which of the M bootstrap replicate posterior distributions Ψ is sampled from. Then, the conditional distribution of Ψ given Z is Ψ *Z*∼ Beta(*α* _*Z*_, *β* _*Z*_), where (*α*_*m*_, *β*_*m*_) are the Beta distribution parameters for the m-th bootstrap replicate’s posterior distribution. The law of total variance states that we can decompose the variance of Ψ into the variance of a conditional expectation and the expectation of a conditional variance. That is:

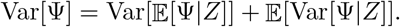

These conditional expectations and variances of a Beta-distributed random variable have closed form:

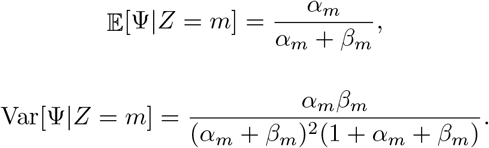

So, the variance of Ψ is:

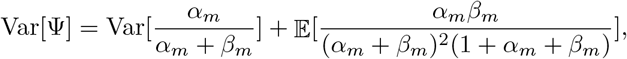

which can be computed by enumerating the means/variances of each component beta distribution and numerically taking their variance/mean. This is faster and correctly calculates the variance of the mixture of bootstrapped posterior distributions on PSI.

### Improved deltapsi prior estimation

MAJIQ V2 calculates the deltapsi prior as a mixture of betas, where mixture components are estimated over deltapsi bins ranging from -1 to 1. MAJIQ V3 updates this procedure in two ways:

1. Doubles the number of bins from 40 to 80.
2. Calculates the mixture variance using expectation maximization, in which the maximization for beta distributions is fitting variance to symmetric beta (rather than doing maximum likelihood, which is not closed form).

This improves the accuracy of the deltapsi prior estimate.

## Supplemental Results

### V2 vs V3 Stratified by Event Type and Coverage

The main text and figure reported results for a “small” evaluation set that included 10 cerebellum and 10 muscle-skeletal RNA-seq experiments from GTEx (see Figure 1). Here we present the same results stratified by event type (retained intron, junction) and total LSV coverage *T* calculated as the sum over junctions. In all results, each LSV quantified by both tools is represented by one junction selected uniformly at random. The thresholds 300 (low vs mid coverage) and 800 (mid vs high coverage) correspond to rounded-off splits of the data into 3 near-equal numbers of events. The PSI results use the V3 PSI quantifications for thresholding. In the DeltaPSI (dPSI) results, only LSVs quantified in the same coverage tier by both tools are included in the comparison.

For each group of LSV quantifications, the Pearson correlation *r* is reported. The lowest-correlated group is the low-coverage retained introns, with *r* = 0.888 for PSI quantifications. Overall, low-coverage groups have slightly lower correlation than middle- and high-coverage groups, and introns have lower-correlation than junctions. Low-coverage retained introns also have the highest percentage (9.8%) of LSVs deviating more than 10% between MAJIQ V2 and V3 in PSI quantifications. The differences can be explained largely as the result of improvements to the quantification algorithms in MAJIQ V3, including significant updates to retained-intron quantification (see main text).

**Table 1:** Percent of LSVs with difference more than 10% between MAJIQ V2 and V3.

| Coverage: | Low | Mid | High |
| --- | --- | --- | --- |
| Junctions PSI | 4.3 | 4.1 | 1.9 |
| Introns PSI | 9.8 | 5.7 | 2.9 |
| Junctions dPSI | 1.9 | 1.6 | 0.7 |
| Introns dPSI | 3.7 | 2.8 | 1.6 |

**Figure 2:**
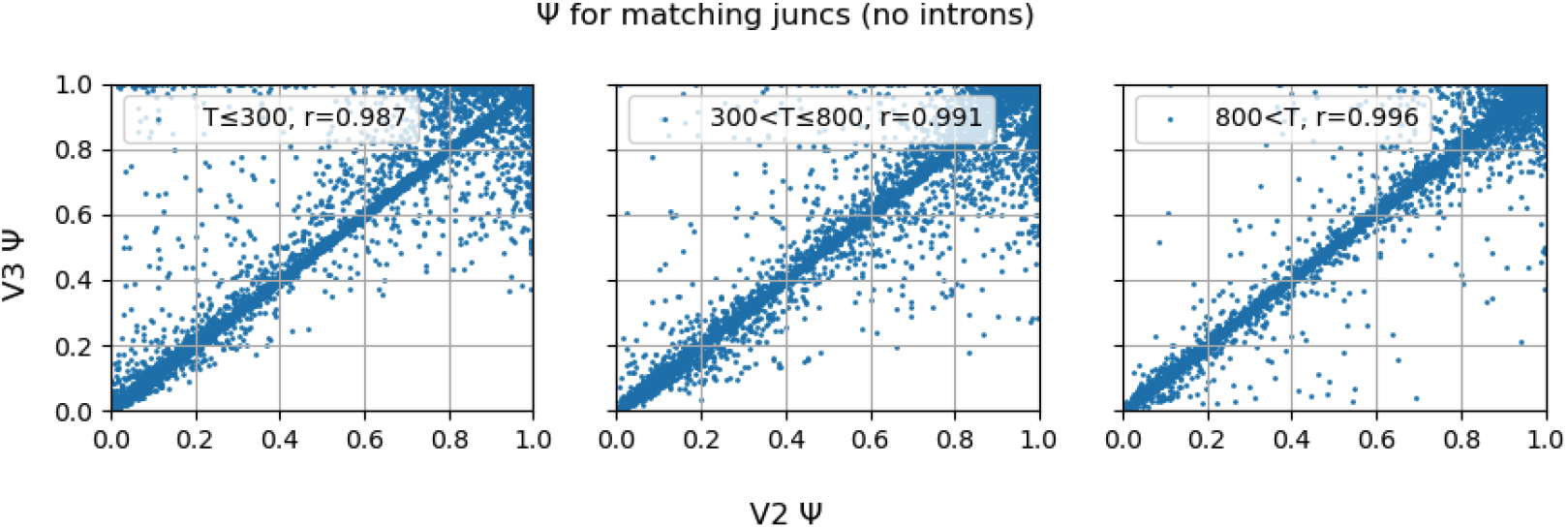
V2 vs V3 PSI for junctions (no retained introns) in 10 cerebellum samples.

**Figure 3:**
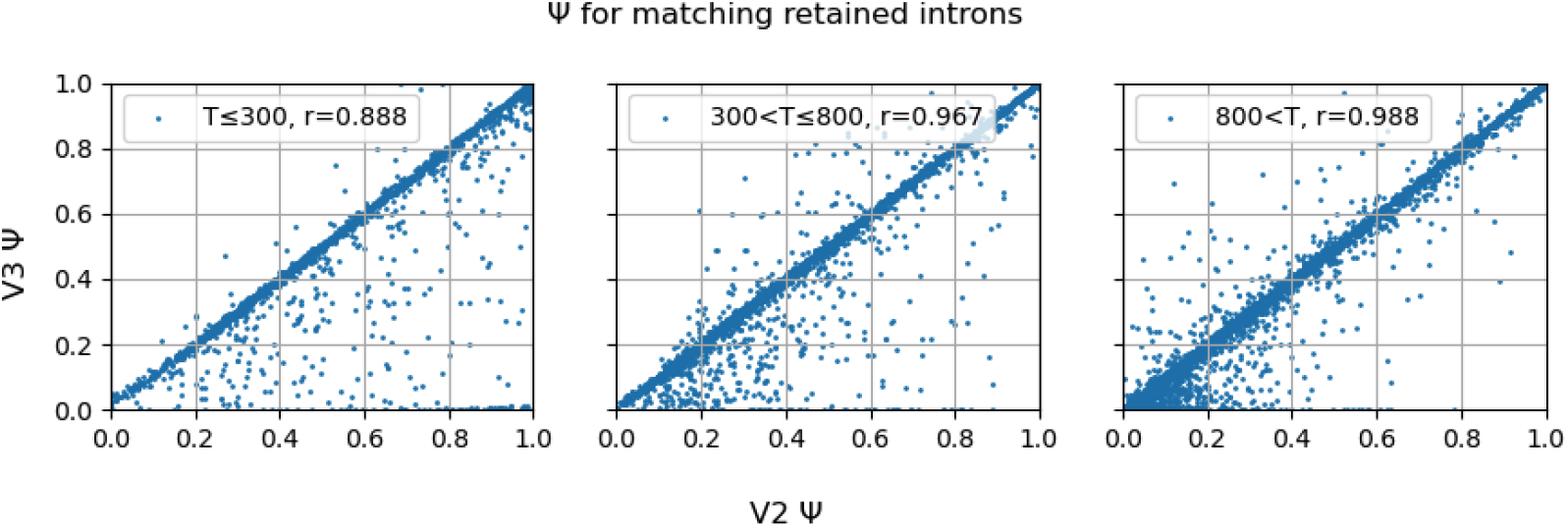
V2 vs V3 PSI for retained introns in 10 cerebellum samples.

**Figure 4:**
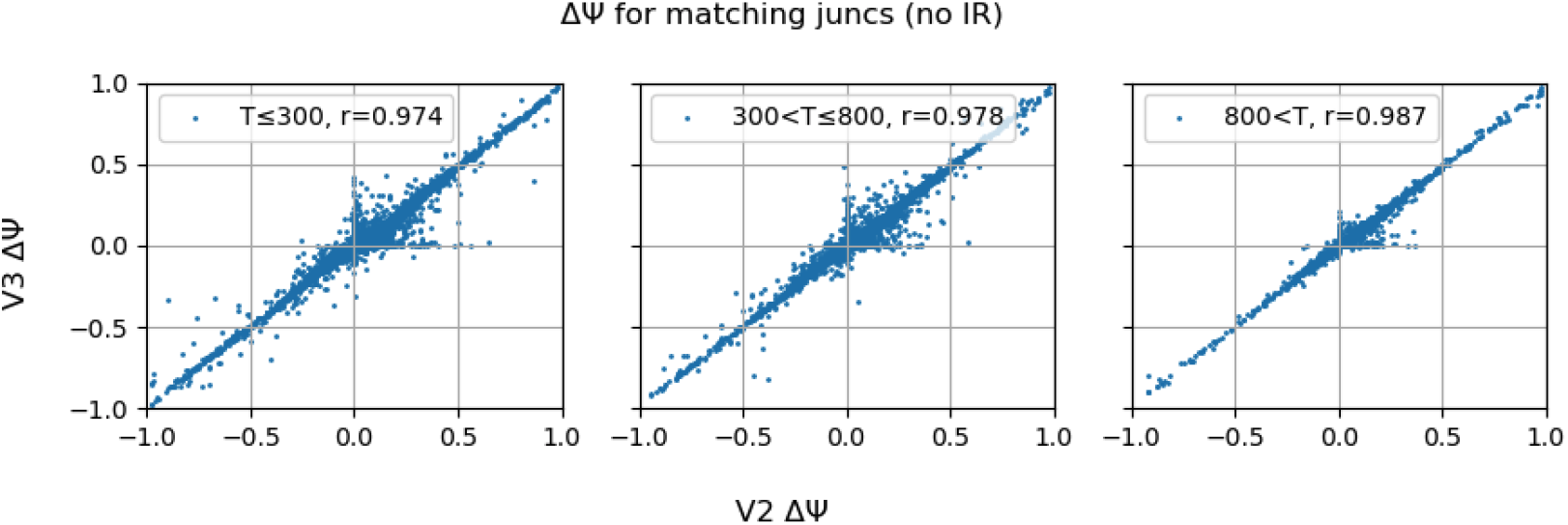
V2 vs V3 DeltaPSI for junctions (no retained introns) between 10 samples each cerebellum and muscle-skeletal.

**Figure 5:**
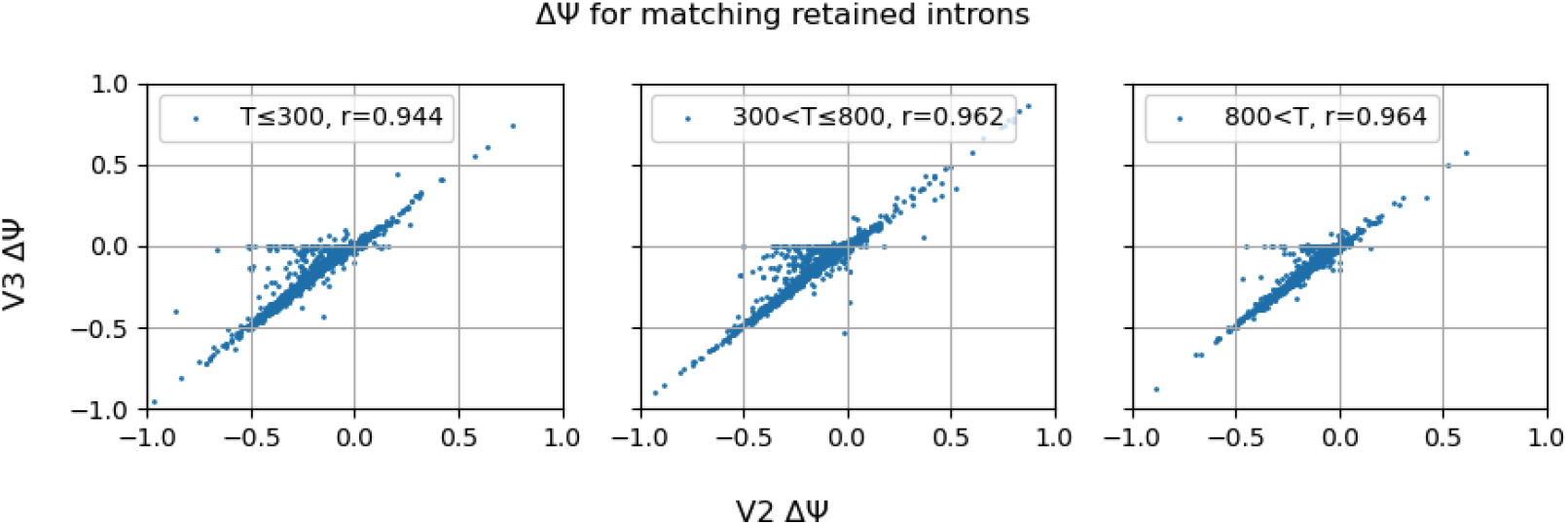
V2 vs V3 DeltaPSI for retained introns between 10 samples each cerebellum and muscle-skeletal.

## References

[1] Qun Pan, Ofer Shai, Leo J. Lee, Brendan J. Frey, and Benjamin J. Blencowe. Deep surveying of alternative splicing complexity in the human transcriptome by high-throughput sequencing. Nature Genetics, 40:1413–1415, 2008. ISSN 1546-1718. doi: 10.1038/ng.259. URL https://www.nature.com/articles/ng.259.

[2] Eric T. Wang, Rickard Sandberg, Shujun Luo, Irina Khrebtukova, Lu Zhang, Christine Mayr, Stephen F. Kingsmore, Gary P. Schroth, and Christopher B. Burge. Alternative isoform regulation in human tissue transcriptomes. Nature, 456:470–476, 2008. ISSN 1476-4687. doi: 10.1038/nature07509. URL https://www.nature.com/articles/nature07509.

[3] Marina M. Scotti and Maurice S. Swanson. Rna mis-splicing in disease. Nature Reviews Genetics, 17:19–32, 2016. ISSN 1471-0064. doi: 10.1038/nrg.2015.3. URL https://www.nature.com/articles/nrg.2015.3.

[4] Diana Baralle and Emanuele Buratti. Rna splicing in human disease and in the clinic. Clinical Science, 131:355–368, 2017. ISSN 1470-8736. doi: 10.1042/CS20160211. URL https://portlandpress.com/clinsci/article-abstract/131/5/355/71590/RNA-splicing-in-human-disease-and-in-the-clinic.

[5] Minghao Jiang, Shiyan Zhang, Hongxin Yin, Zhiyi Zhuo, and Guoyu Meng. A comprehensive benchmarking of differential splicing tools for rna-seq analysis at the event level. Briefings in Bioinformatics, 24, 2023. ISSN 1477-4054. doi: 10.1093/bib/bbad121. URL https://academic.oup.com/bib/article/24/3/bbad121/7108868.

[6] Jorge Vaquero-Garcia, Alejandro Barrera, Matthew R. Gazzara, Juan Gonzalez-Vallinas, Nicholas F. Lahens, John B. Hogenesch, Kristen W. Lynch, and Yoseph Barash. A new view of transcriptome complexity and regulation through the lens of local splicing variations. eLife, 2016. ISSN 2050-084X. doi: 10.7554/eLife.11752. URL https://elifesciences.org/articles/11752.

[7] Jorge Vaquero-Garcia, Joseph K. Aicher, San Jewell, Matthew R. Gazzara, Caleb M. Radens, Anupama Jha, Scott S. Norton, Nicholas F. Lahens, Gregory R. Grant, and Yoseph Barash. Rna splicing analysis using heterogeneous and large rna-seq datasets. Nature Communications, 14, 2023. ISSN 2041-1723. doi: 10.1038/s41467-023-36585-y. URL https://www.nature.com/articles/s41467-023-36585-y.

[8] Seong Woo Han, San Jewell, Andrei Thomas-Tikhonenko, and Yoseph Barash. Contrasting and combining transcriptome complexity captured by short and long rna sequencing reads. BioRxiv [preprint], 2023. doi: 10.1101/2023.11.21.568046. URL https://www.biorxiv.org/content/10.1101/2023.11.21.568046v1.

[9] Barry Slaff, Caleb M. Radens, Paul Jewell, Anupama Jha, Nicholas F. Lahens, Gregory R. Grant, Andrei Thomas-Tikhonenko, Kristen W. Lynch, and Yoseph Barash. Moccasin: a method for correcting for known and unknown confounders in rna splicing analysis. Nature Communications, 12, 2021. ISSN 2041-1723. doi: 10.1038/s41467-021-23608-9. URL https://www.nature.com/articles/s41467-021-23608-9.

[10] John Lonsdale, Jeffrey Thomas, and Mike Salvatore et al. The genotype-tissue expression (gtex) project. Nature Genetics, 45:580–585, 2013. ISSN 1470-8736. doi: 10.1038/ng.2653. URL https://www.nature.com/articles/ng.2653.

